# s-aligner: a greedy algorithm for non-greedy de novo genome assembly

**DOI:** 10.1101/2021.02.02.429443

**Authors:** Juanjo Bermúdez

## Abstract

Genome assembly is a fundamental tool for biological research. Particularly, in microbiology, where budgets per sample are often scarce, it can make the difference between an inconclusive result and a fully valid conclusion. Identifying new strains or estimating the relative abundance of quasi-species in a sample are some example tasks that can’t be properly accomplished without previously generating assemblies with little structure ambiguity and covering most of the genome. In this work, we present a new genome assembly tool based on a greedy strategy. We compare the results obtained applying this tool to the results obtained with previously existing software. We find that, when applied to viral studies, comparatively, the software we developed often gets far larger contigs and higher genome fraction coverage than previous software. We also find a significant advantage when applied to exceptionally large virus genomes.

## Introduction

Some months ago, I posted a paper to a preprint server. I made public that work with the only intention to support another one. I thought I didn’t want to explain that particular method in the same paper as my main finding, but I considered that this method should be available anyway somehow to whoever wanted to dig deeper into my research. So, I wrote a separate paper explaining that method. Given that I didn’t care much if anybody was going to consider it seriously, and from previous experiences in a public online forum indicating that it wouldn’t be, I decided to use an overly honest tone, admitting the naivety of its conception and its limited partial utility over its initial intended utility. I thought nobody would attack it if I previously, humbly, admitted the limitations of its conception myself.

It was attacked. And in so much intensity that as a side effect it became one of the top 5% research outputs scored at Altmetric. 97th percentile for outputs of the same age. 98th percentile for outputs of the same age and source. Nobody ever has cited it, though, and nobody has shown interest to test the software even if I took the trouble to develop a free online site so that anybody could. Some voices even claimed that my work should be proposed to an Ig Nobel prize.

Now I present a new software that could not have been developed without applying that method (1). It could not have been developed using BLAST (2) for finding local alignments. BLAST is not well-suited for finding local alignments of short sequences, as it was exposed in my mocked paper, even if overall it might be a far better tool for finding local alignments.

And that’s the philosophy that took me to develop some other basic algorithms that overall do not present general advantages over the commonly established ones, but that in some particular circumstances, they do. When put together, all these algorithms let me experiment with new strategies for genome assembly.

Fruit of that experimentation I present here an algorithm with significant advantages over other known software for viral genome assembly.

## Method

There are two established strategies for de novo genome assembly.

- Greedy algorithm assemblers are assemblers that find local optima in alignments of smaller reads. Greedy algorithm assemblers typically feature several steps: 1) pairwise distance calculation of reads, 2) clustering of reads with the greatest overlap, 3) assembly of overlapping reads into larger contigs, and 4) repeat. These algorithms are claimed not to work well for larger read sets, not easily reaching a global optimum in the assembly, and not performing well on read sets that contain repeat regions.
- Graph method assemblers come in two varieties: string and De Bruijn. These methods are claimed as an important step forward because of reaching a global optimum instead of a local optimum The De Bruijn graph method has become the most popular in the age of next-generation sequencing.

The algorithm presented in this work makes use of a greedy algorithm strategy but not exactly the Overlap-Layout-Consensus method used by most algorithms in the past.

### A. Overlap

This step is similar to the one in any overlap-layout-consensus algorithm but in this case, a local-alignment search tool is applied to find the overlaps. The requirements for this tool are i) being fast enough, ii) being accurate enough for aligning short sequences, and iii) not being too resource-greedy so that it can be run on a standard desktop computer.These requirements are covered by using a tool like the one described in SLAST (1).

### B. Layout

This is the step in which this algorithm differs the most from a typical overlap-layout-consensus algorithm.

Instead of looking for a Hamiltonian path in a graph connecting overlapped reads, an alternative simplistic approach is applied.

The method is as follows:

**Algorithm 1.**
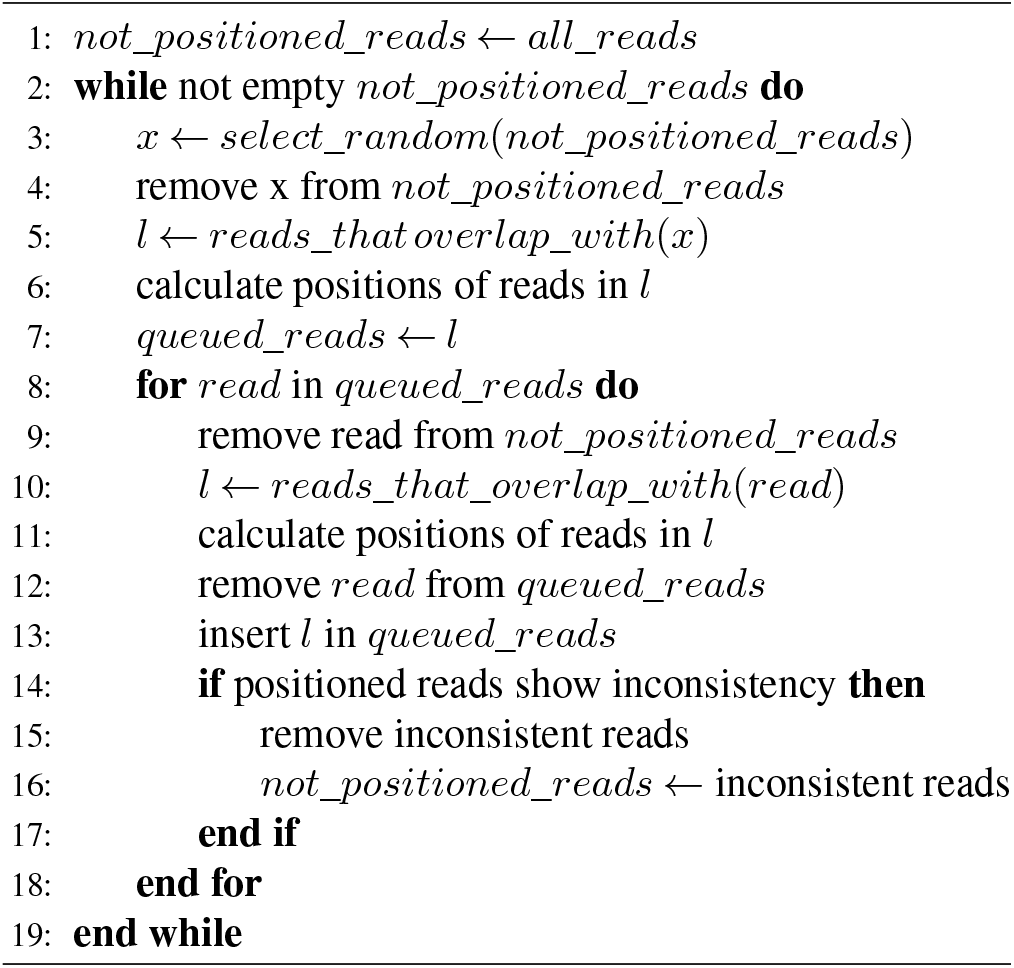
Overlap-Layout

Or explained in plain text:

1. Selecting a random read
2. Calculating the position of all reads that overlap with that read.
3. Repeating with every read that we added.

a. Every while, take a look at how the reads look once positioned. If the reads seem inconsistent due to the presence of artifacts or a divergent path caused by a repeated sequence, identify the reads provoking the inconsistency, remove them, and continue. See figure 1
4. If there are remaining reads that have not been aligned yet (because they didn’t overlap any read previously assembled or because they were removed due to inconsistency), select a random read from the ones not assembled yet and start again. Every concatenation of reads departing from a different read will be considered a contig.

**Fig. 1.**
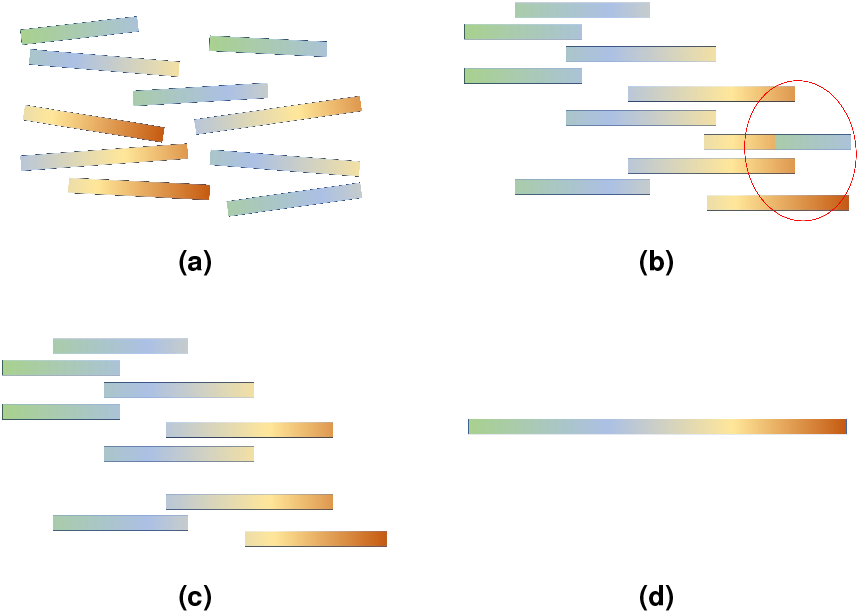
(a) A set of reads. (b) The reads aligned including a chimeric one. (c) Removal of the chimeric read. (d) The final contig.

**Fig. 2.**
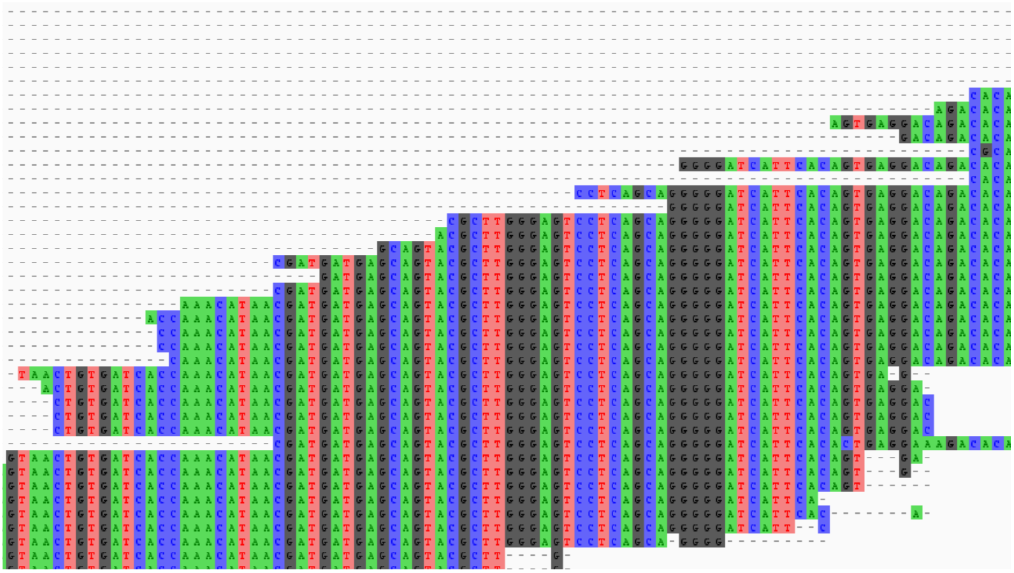
Some reads aligned to form a contig visualized with AliView (3)

### C. Consensus

This step, again, is similar to the one in any overlap-layout-consensus algorithm, but in this case, extra precision is sought by making a multiple-alignment of the reads after being positioned.

Figure 3 shows the usual aspect of one of these alignments.

**Fig. 3.**
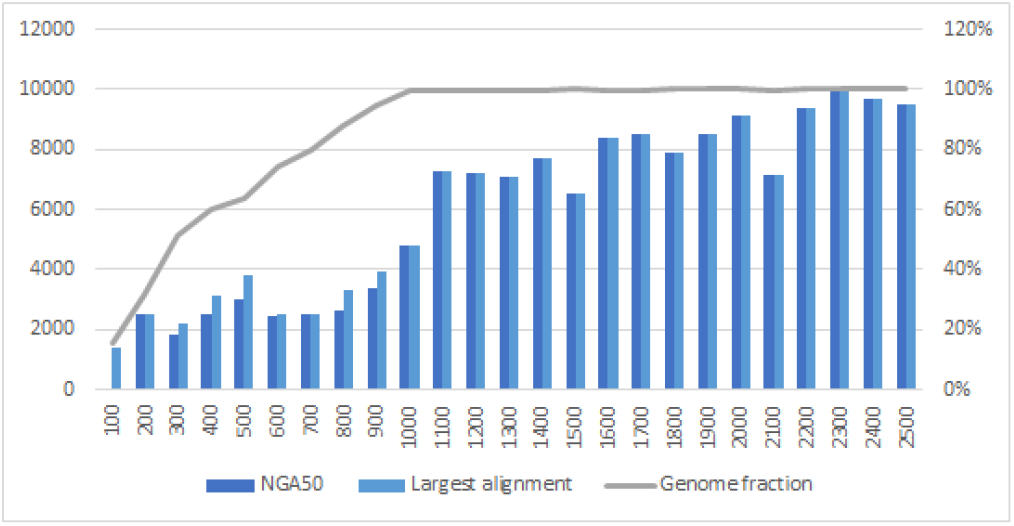
s-aligner metrics for HIV-labmix (20.000x) set in intervals of 100 reads processed

## Characteristics

### A. Interactive

Like other greedy algorithms, s-aligner can be scheduled according to different strategies and you don’t need to wait for it to complete to see how it is performing: you can monitor and modify your strategy interactively. For example, you can select at every step how many seed-reads to randomly select to start finding overlaps. If you start with one read it’s more likely that you get a large contig in which it is included, and in contrast, if you start with many it’s more likely that you end up with shorter contigs but covering different strains or different regions of the genome. You can also select for how long to continue assembling until a result is generated to be evaluated. Once evaluated, you can continue assembling from that assembly by just selecting the assembly id.

### B. Quality/Speed adjusting

Another way in which you can configure s-aligner is by selecting how fast it tries to assemble. The faster it goes, the fewer reads it uses to cover every nucleotide position. That means that going faster more errors might be included in the final sequence. On the other hand, by going faster you can get results earlier and adjust your strategy for a posterior assembly using a more accurate approach.

### C. No need for paired-end reads

Another peculiarity of s-aligner is that it does not make use, for now, of pairing information between reads. If you have two separated files for forward and reverse paired reads, you can omit one of them, and if there is enough data your result will be equally reliable. Think that s-aligner will only use a small fraction of all the available reads when the coverage is high enough. Usually, any set granting coverage over 6x for every nucleotide will be good enough for assembling it.

### D. No quality measurements for input data but for output data

S-aligner does not require quality measurements for the input data. Its performance is exactly the same whether or not such data is available. It simply converts FASTQ inputs to FASTA. The reason? I just didn’t find any way to take advantage of that information and its performance is good enough without finding that. In contrast, you can switch a parameter to make it generate quality measurements for the output: it can generate FASTQ files for the assemblies. The quality measurements are obtained from the consensus step during the assembly. You can also generate files in which all reads used for every contig are aligned. This way you can inspect visually how any particular area of the assembly was obtained. That is sometimes very useful, for example, for determining if a possible mutation is real or just a side-effect of the assembly.

## Benchmarks

The algorithm has been tested with different publicly available benchmark sets and its results have been compared with results for these data sets in peer-reviewed literature.

### A. Claimed gold standard for genome assembly of viral quasispecies

This is a HIV-1 full-length benchmarking data set for haplotype reconstruction methods, sequenced with Illumina MiSeq and 454/Roche GSJunior. Five well-studied HIV-1 strains (HXB2, 89.6, JR-CSF, NL4-3, and YU-2) have been mixed and sequenced. This data set was obtained from Giallonardo et al. (4).

The results from the independent study Baaijens et al. (5) have been collected and compared to the results obtained by s-aligner.

The metrics analyzed are NG50 and Genome Fraction, as these are the most useful metrics for evaluating the potential of generating better scaffolds. QUAST (6) was used for obtaining the metrics from the s-aligner results. Some metrics, like Percent of Mismatches, have been discarded as QUAST doesn’t seem to calculate these correctly for assemblies with Duplication Ratio over 1. Also, these error rates are not reliable metrics to evaluate the assembly as we are not comparing against the real sequence but just a reference. In the case of highly mutable viruses like HIV that makes even less sense.

The results can be seen in table 1.

**Table 1.** HIV labmix

|  | NG50 | Genome fraction |
| --- | --- | --- |
| SAVAGE | 588 | 92,6 |
| SGA | 635 | 32,4 |
| SOAPdenovo2 | 591 | 41,9 |
| SPAdes | 591 | 42,6 |
| metaSPAdes | 3266 | 53,7 |
| s-aligner (Illumina data) | 9641 | 100 |
| s-aligner (Roche data) | 9062 | 100 |

HIV-labmix sample assembly metrics for different assemblers.

Note that s-aligner made no use of paired-end information, therefore a data set generated without paired-end reads would perform exactly the same, while the other assemblers would get even worse results. Single-read sequencing is cheaper, therefore s-aligner could reduce costs of sequencing while increasing the quality of the results.

Apart from measuring the overall assembly performance, it was also analyzed what was the performance for every strain in the sample. The results were similar, always getting NG50 over 9000. Therefore, apart from improving assembly results s-aligner also demonstrated good qualities for identifying and extracting strains in mixed samples. The composition of the sample can be seen in table 3 and the results for every strain in table 2.

**Table 2.** HIV labmix strains

|  | NG50 | Genome fraction |
| --- | --- | --- |
| YU-2 strain | 9551 | 100 |
| 89.6 strain | 9244 | 99,94 |
| NL4-3 strain | 9641 | 100 |
| JR-CSF strain | 9546 | 100 |
| HXB2 strain | 9641 | 100 |

Metrics for s-aligner assembly applied to different strains present in the HIV-labmix sample.

**Table 3.** Characteristics of benchmarking data sets

| Data set | Virus type | Genome length (bp) | Average coverage | Strain count | Strain abundance | Pairwise divergence |
| --- | --- | --- | --- | --- | --- | --- |
| 600x HIV mix | HIV-1 | 9478-9719 | 600x | 5 | 20% | 1%-6% |
| 5-strain HIV mix | HIV-1 | 9478-9719 | 20.000x | 5 | 5%-28% | 1%-6% |
| 10-strain HCV mix | HCV-1a | 9273-9311 | 20.000x | 10 | 5%-19% | 6%-9% |
| 3-strain ZIKV mix | ZIKV | 10251-10269 | 20.000x | 3 | 16%-60% | 3%-10% |
| 15-strain ZIKV mix | ZIKV | 10251-10269 | 20.000x | 15 | 2%-13% | 1%-10% |
| Lab mix | HIV-1 | 9478-9719 | 20.000x | 5 | 10%-30% | 1%-6% |

For each benchmark, we specify virus type, genome length, average coverage, strain count, relative abundance, and pairwise divergence. For the 600x HIV mix, the strains were homogeneously distributed with a relative abundance of 20% each.

### B. De novo assembly of simulated viral samples

Another benchmark set in the public literature corresponds to the one obtained by simulation of viral samples in Baaijens et al. (5). They describe that data set in the following way. “*We created five simulated data sets for benchmarking, consisting of 2 × 250-bp Illumina MiSeq reads and representing quasispecies infections from different viruses: human immunodeficiency virus (HIV), hepatitis C virus (HCV), and Zika virus (ZIKV). We varied the number of strains per sample as well as the relative abundances of those strains and the pairwise divergence between strains. To get data sets as realistic as possible, we used true viral genomes from the NCBI database and Illumina MiSeq error profiles during simulations*.”

The results for that benchmark set are in tables 4, 5, 6, and 7.

**Table 4.**
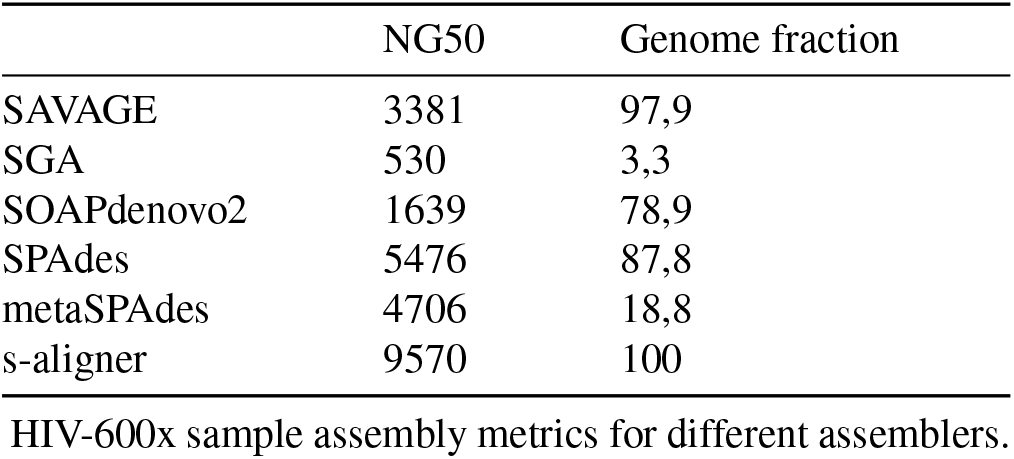
HIV-600x

|  | NG50 | Genome fraction |
| --- | --- | --- |
| SAVAGE | 3381 | 97,9 |
| SGA | 530 | 3,3 |
| SOAPdenovo2 | 1639 | 78,9 |
| SPAdes | 5476 | 87,8 |
| metaSPAdes | 4706 | 18,8 |
| s-aligner | 9570 | 100 |

HIV-600x sample assembly metrics for different assemblers.

**Table 5.** HIV-20000x

|  | NG50 | Genome fraction |
| --- | --- | --- |
| SAVAGE | 4913 | 92,6 |
| SGA | 650 | 32,4 |
| SOAPdenovo2 | 516 | 41,9 |
| SPAdes | 5873 | 42,6 |
| metaSPAdes | 5159 | 53,7 |
| s-aligner | 9916 | 100 |

HIV-20000x sample assembly metrics for different assemblers.

**Table 6.** HCV-20000x

|  | NG50 | Genome fraction |
| --- | --- | --- |
| SAVAGE | 8248 | 99,6 |
| SGA | 658 | 18,1 |
| SOAPdenovo2 | 531 | 22,0 |
| SPAdes | 8582 | 91,3 |
| metaSPAdes | 1549 | 45,9 |
| s-aligner | 9285 | 96,3 |

HCV-20000x sample assembly metrics for different assemblers.

**Table 7.** ZIKV-20000x

|  | NG50 | Genome fraction |
| --- | --- | --- |
| SAVAGE | 2103 | 99,4 |
| SGA | 0 | 0 |
| SOAPdenovo2 | 562 | 21,0 |
| SPAdes | 2577 | 65,6 |
| metaSPAdes | 3926 | 17,5 |
| s-aligner | 10161 | 95,0 |

ZIKV-20000x sample assembly metrics for different assemblers.

#### B.1. Hardware Requirements

Table 8 shows some hardware performance metrics found while benchmarking s-aligner.

**Table 8.**
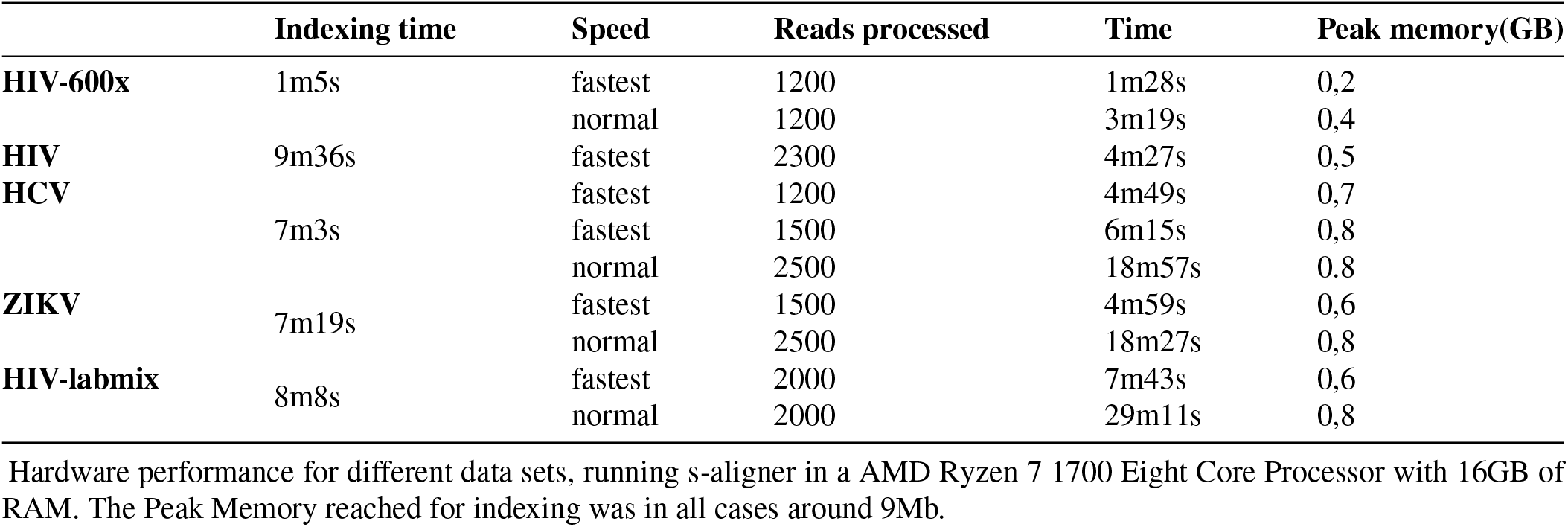
Hardware performance

Note that these results correspond to the application of standard parameters. The speed of assembly could likely be increased, for example, by indexing a smaller subset of reads. Around 200.000 reads per run (when available) were selected for indexing on all tests previously shown.

All results whose timing have been measured in table 8 had similar NG50 to s-aligner results in previous tables, but note that you can get results with lower NG50 by processing fewer reads, therefore spending less time, and yet these would be improved results over the ones from other assemblers.

### C. Multiple-strain samples of large DNA viruses

S-aligner has also been tested with phages and other large viruses. It also performs well for viruses under 150 kbp. In some cases, though, like the one in the following benchmark set (viruses around 250 kbp), it does not perform well enough by itself, but it does if an additional assembly step is performed. S-aligner usually generates more contigs than other assemblers and many of these are large sequences that indeed overlap with other contigs. If these contigs are assembled in a second step, you finally get even larger contigs. And given that these contigs are usually already quite large, the best way to assemble these is using a genome assembler tool for large reads, like Flye (7) or Canu (8).

See in figure 4 how s-aligner contigs usually look when aligned to a reference genome. Note the overlapping

**Fig. 4.**
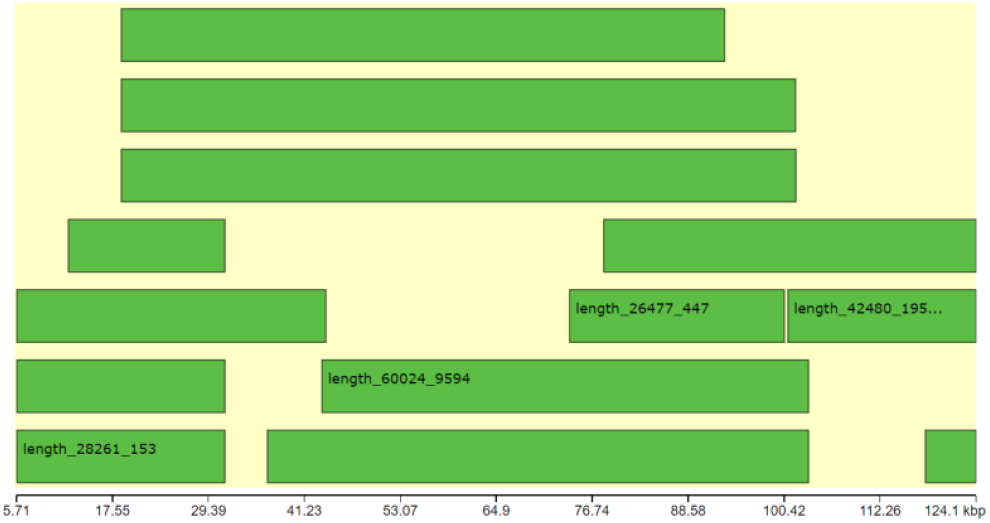
Capture of QUAST/Icarus (6) (9) chart showing contigs aligned to a reference genome.

Therefore, to assemble the following benchmark set a two-step strategy was employed:

1. Use s-aligner with the raw reads as input.
2. Use Flye with the contigs obtained by s-aligner as input. Optionally, merge contigs obtained at this step with the contigs obtained from s-aligner.

The following benchmark set as well as the results for the other genome assemblers have been extracted from the following independent report: Deng et al. (10).

They created in vitro viral strain mixtures mimicking clinical samples from patients with mixed strain infections. For this, they combined viral DNA of the HCMV strains TB40/E BAC4 and AD169 (designated as “TA”), derived directly from bacterial artificial chromosomes (BAC) with these viral genomes and prepared from Escherichia coli, or the strains TB40/E BAC4 and Merlin (designated as “TM”), which were amplified in human cell-cultures, respectively, at mixing ratios of 1:1, 1:10 and 1:50. The NG50 results for this benchmark are in figure 5.

**Fig. 5.**
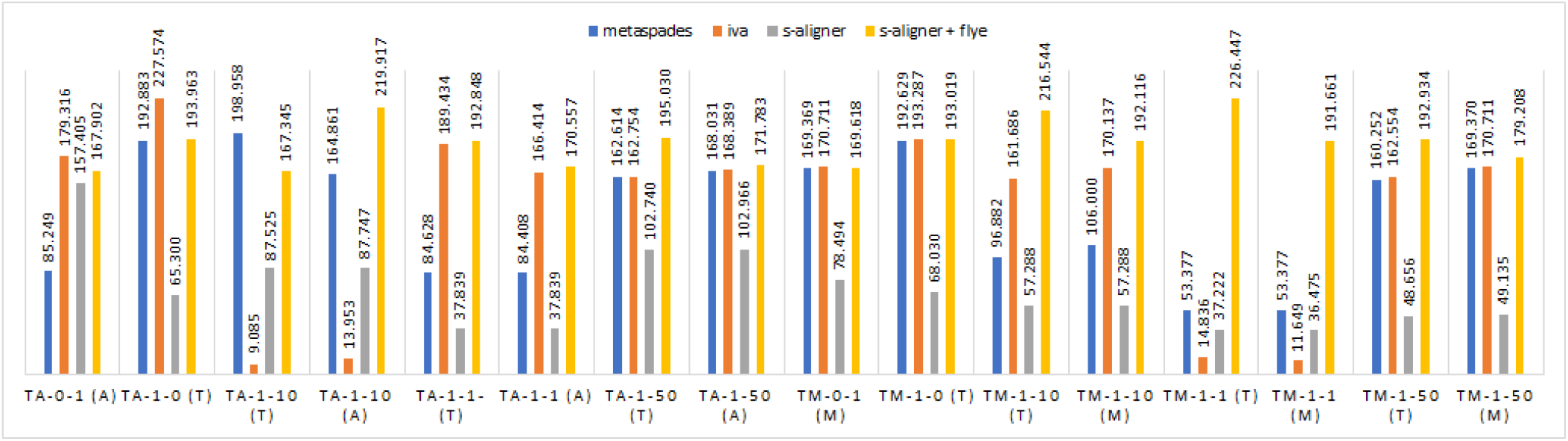
NG50 metric for results in the large DNA virus set. Between brackets, the strain making as reference for the measurement.

We can see how, also with larger genomes, we can obtain better assemblies if applying s-aligner at some step. We see how this method also outperforms assemblers designed for larger genomes, like SPAdes or metaSPAdes (assemblers with lower performances in the original study have been omitted in the figure). In this case, it gets in average a 64% increase of performance for the NG50 metric. The genome Fraction, in this case, was on par with the second-best assembler, and near to 100% on average. Please note, in addition, that these are not the best possible results that you can get with s-aligner, but just some results good enough that were obtained in a reasonable time. You could improve the results by fitting your strategy. Also note that if you usually get your data using a similar protocol, you will be likely able to establish an strategy that better fits with the characteristics of your data.

Other assemblers evaluated in the original study from which the data set was obtained (10) but whose results have not been included in this study due to their significant lower performance with this data set are: SPAdes (11), Megahit (12), ABySS (13), Ray (14), IDBA-UD (15), Tadpole (16), Vicuna (17) and Savage (5).

## Conclusions

The solution reported in this document has demonstrated better performance for assembling genomes of small size than any other genome assembler previously benchmarked in the public peer-reviewed literature. This increased performance is on average 110% for short virus genomes and 64% for exceptionally large virus genomes. The increased performance has also been confirmed on many data sets found in the SRA database.

It performs well both with simulated and real data, with data from different hardware providers, and with or without paired-end information. It even performs better than other software when using largely poorer data than other assemblers (600x vs 20.000x).

Overall, all these advantages mean that in a crisis like the one caused by COVID-19, the hundreds of thousands of sequencings being done around the world could make use of cheaper resources to obtain equivalent or superior quality in the results. That could have a significant impact on the management of the crisis.

For all these reasons (its decreased dependence on resources and reduction of costs) I ended up titling this document “s-aligner: a greedy algorithm for non-greedy genome assembly”

## Discussion

We demonstrate how sometimes the best solution to a problem is not found by having an encyclopedic knowledge of the available methods in the state of the art but just by naively developing intuitive solutions ex professo, even if that is not what the academic life promotes. No advanced technical method which wasn’t already available 50 years ago was required for developing this solution, and yet it wasn’t until now. That might be suggesting, in my opinion, inefficiencies in the way how actual science and academic organizations promote and recognize talent. And this also inevitably brings the question of how many fields in science may also be missing quasi-obvious solutions due to ill incentives in the academic life.

This problem is increased by other already detected problems in the bioinformatics field (18).

More work in this direction is needed now to adapt this algorithm to other genome assembly protocols of equal or bigger relevance, like transcriptome de novo assembly or genome assembly for larger genomes.

## Supporting information

assembly results

## Data availability statement

The data underlying this article are available in the article and in its online supplementary material.

Benchmark sets employed for this study were made available by the original authors of the previous study at https://bitbucket.org/jbaaijens/savage-benchmarks

The sequence data for the large-dna study were deposited by the authors in ENA with accession number PRJEB32127.

The s-aligner software is available for free at https://contignant.com for first-time users. It’s free to use for 15 days after installation. No personal identification is required but a contact email must actually be provided for downloading it. Contact me if, for any special circumstance, you need an extension of the trial period for evaluating the results in this document.

## Competing interests

There is NO Competing Interest.

## Notes

https://contignant.com

### Competing Interest Statement

The authors have declared no competing interest.

